# Labkit: Labeling and Segmentation Toolkit for Big Image Data

**DOI:** 10.1101/2021.10.14.464362

**Authors:** Matthias Arzt, Joran Deschamps, Christopher Schmied, Tobias Pietzsch, Deborah Schmidt, Robert Haase, Florian Jug

## Abstract

We present Labkit, a user-friendly Fiji plugin for the segmentation of microscopy image data. It offers easy to use manual and automated image segmentation routines that can be rapidly applied to single- and multi-channel images as well as to timelapse movies in 2D or 3D. Labkit is specifically designed to work efficiently on big image data and enables users of consumer laptops to conveniently work with multiple-terabyte images. This efficiency is achieved by using ImgLib2 and BigDataViewer as the foundation of our software. Furthermore, memory efficient and fast random forest based pixel classification inspired by the Waikato Environment for Knowledge Analysis (Weka) is implemented. Optionally we harness the power of graphics processing units (GPU) to gain additional runtime performance. Labkit is easy to install on virtually all laptops and workstations. Additionally, Labkit is compatible with high performance computing (HPC) clusters for distributed processing of big image data. The ability to use pixel classifiers trained in Labkit via the ImageJ macro language enables our users to integrate this functionality as a processing step in automated image processing workflows. Last but not least, Labkit comes with rich online resources such as tutorials and examples that will help users to familiarize themselves with available features and how to best use Labkit in a number of practical real-world use-cases.

## Introduction

In recent years, new and powerful microscopy and sample preparation techniques have emerged, such as lightsheet (1), super-resolution microscopy (2–6), modern tissue clearing (7, 8), or serial section scanning electron microscopy (9, 10) enabling researchers to observe biological tissues and their underlying cellular and molecular composition and dynamics in unprecedented details. To localize objects of interest and exploit such rich datasets quantitatively, scientists need to perform image segmentation, *e*.*g*. dividing all pixels in an image into foreground pixels (part of objects of interest) and background pixels.

The result of such a pixel classification is a binary mask, or a (multi-)label image if more than one foreground class are needed to discriminate different objects. Masks or label images enable downstream analysis that extract biologically meaningful semantic quantities, such as the number of objects in the data, morphological properties of these objects (shape, size, *etc*.), or tracks of object movements over time. In most practical applications, image segmentation is not an easy task to solve. It is often rendered difficult by the sample’s biological variability, imperfect imaging conditions (*e*.*g*. leading to noise, blur, or other distortions), or simply by the complicated three-dimensional shape of the objects of interest.

Current research in bio-image segmentation focuses primarily on developing new deep learning approaches, with more classical methods currently receiving little attention. Algorithms, such as StarDist (11), DenoiSeg (12), PatchPerPix (13), PlantSeg (14), CellPose (15), or EmbedSeg (16) have continuously raised the state-of-the art and outperform classical methods in quality and accuracy of achieved automated segmentation. While these approaches are very powerful indeed, deep learning does require some expert knowledge, dedicated computational resources not everybody has access to, and typically large quantities of densely annotated ground-truth data to train on.

More classical approaches, on the other hand, can also yield results that enable the required analysis, while often remaining fast and easy to use on any laptop or workstation. Examples for such methods range from intensity thresholding and seeded watershed, to shallow machine learning approaches on manually chosen or designed features. One crucial property of shallow techniques, such as random forests (17), is that they require orders of magnitude less ground-truth training data than deep learning based methods. Hence, multiple software tools pair them with user-friendly interfaces, *e*.*g*. CellProfiler (18), Ilastik (19), QuPath (20), and Trainable Weka Segmentation (21). The latter specializes in random forest classification and is available within Fiji (22), a widely-used image analysis and processing platform based on ImageJ (23) and ImageJ2 (24). It is, regrettably, not capable of processing very large datasets due to its excessive demand for CPU memory, leaving the sizable Fiji community with a lack of user-friendly pixel classification or segmentation tools that can operate on large multi-dimensional data.

The required foundations for such a software tool have in recent years been built by the vibrant research software engineering community around Fiji and ImageJ2. The problem of handling large multi-dimensional images has been addressed by a generic and powerful library called ImgLib2 (25). Additionally, a fast, memory-efficient, and extensible image viewer, the BigDataViewer (26), enables tool developers to create intuitive and fast data handling interfaces.

Here, we present an image labeling and segmentation tool called Labkit. It combines the power of ImgLib2 and BigDataViewer with a new implementation of random forest pixel classification. Labkit features a user-friendly interface allowing for rapid scribble labeling, training, and interactive curation of the segmented image. Labkit also allows users to fully manually label pixels or voxels in the loaded images. It can be easily installed in Fiji via its updater, and it can directly be called from Fiji’s macro programming language. Labkit additionally features GPU acceleration using CLIJ (27), and can be used on high performance computing (HPC) clusters thanks to a command-line interface.

### Image Segmentation with Labkit

Labkit’s user interface is built around the BigDataViewer (26), which allows interactive exploration of image volumes of any size and dimension on consumer computing hardware (**Fig. 1A, B**). Beyond the common BigDataViewer features, users have access to a set of simple drawing tools to manually paint or correct existing labels on image pixels in 2D and voxels in 3D. Importantly, the raw data is never modified by any such actions. Pixel and voxel labels are grouped by classes in individual layers. Each class is represented by a modifiable color, and can be used to annotate different types of objects and structures of interest in the image.

**Fig. 1.**
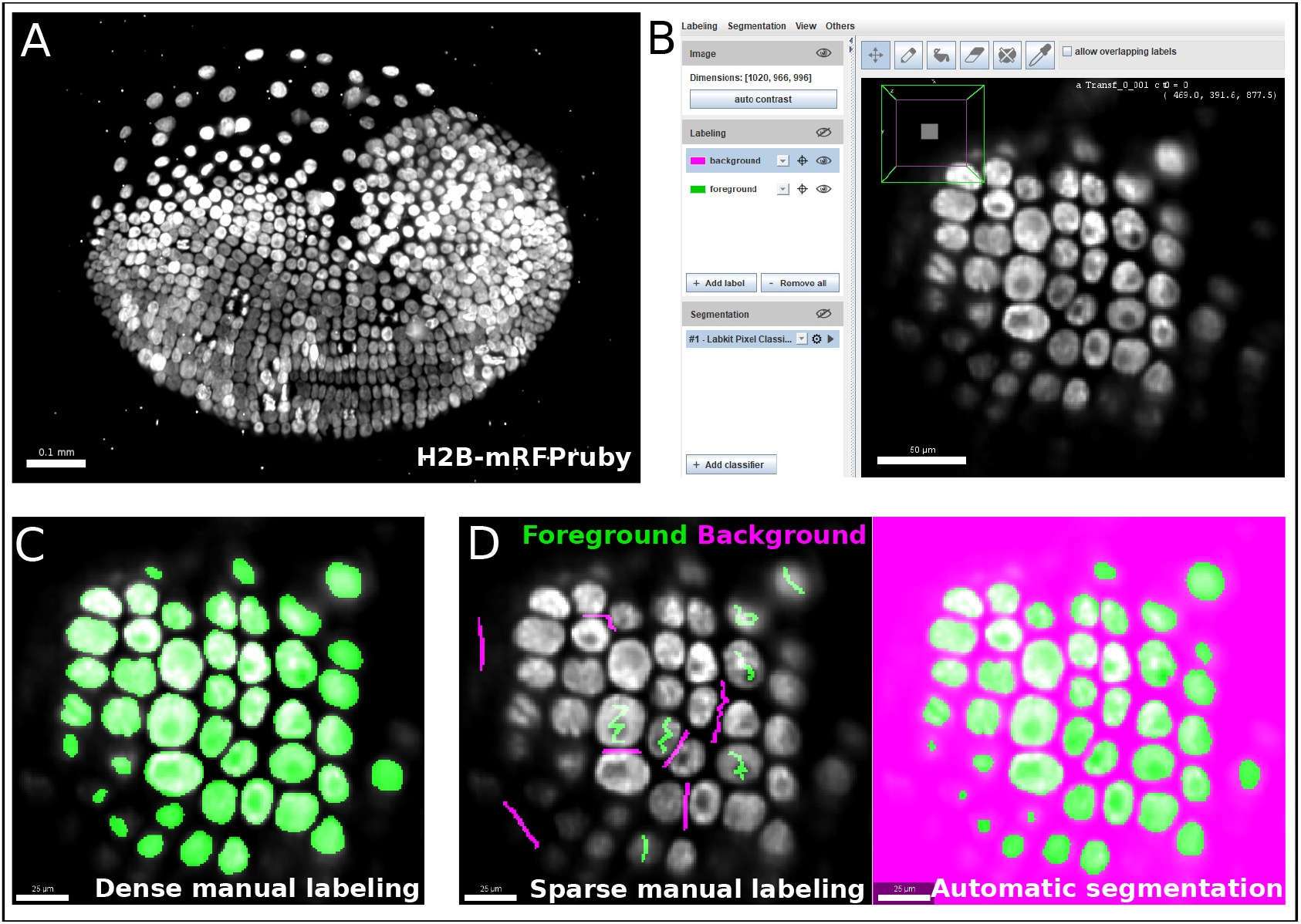
Labkit allows easy manual labeling and automatic segmentation of large image volumes: **(A)** Maximum intensity projection of a single time point from a ∼1 TB timelapse of a developing *Parhyale* embryo imaged live with lightsheet microscopy. **(B)** Labkit’s user interface is based on BigDataViewer and allows visualizing and interacting with large volumes of image data. A slice of the developing *Parhyale* embryo is shown. **(C)** Users can label large datasets with dense manual annotations using Labkit’s drawing interface. **(D)** A core feature of Labkit is the rapid segmentation of large image data using sparse manual labels (scribbles) combined with random forest pixel classification to automatically produce the final segmentation. Scale bars 100 μm **(A)**, 50 μm **(B)**, 25 μm **(C, D)**,

Thanks to the intuitive interface design, users can efficiently segment their images by manually drawing dense labels on the entire image (**Fig. 1C**). Labels that are generated with the drawing tools can directly be saved as images or exported to Fiji for downstream processing. Dense manual labelings of complete images or volumes created with Labkit can be used to manually segment objects, as was done previously to mask particles in cryo-electron tomograms of *Chlamydomonas* (28).

However, this process is very time consuming and doesn’t scale well to large data. Labkit is therefore often used to densely and manually label a subset of the image data, which is then used as ground-truth for supervised deep learning approaches. Published examples include the generation of ground-truth training data for a mouse and a *Platyneris* dataset in order to segment cell nuclei with EmbedSeg (16). Labkit is also suggested as a tool of choice for ground-truth generation by other deep learning methods (11, 12, 29). Still, manually generating sufficient amount of ground-truth training labels for existing deep learning methods remains a cumbersome and tedious task.

In order to create high quality segmentation while maintaining a manageable amount of user input, a core feature of Labkit is a random forest based pixel classification (17) based on Weka (30, 31), newly implemented and optimized for speed. When using this feature, instead of annotating entire objects, a random forest is trained on only a few pixel annotations per class. Such sparse manual labels, or scribbles (see **Fig. 1D**, left), are directly drawn by users over the image. Naturally, the sparse labels must be drawn on correct and representative pixels from each pixel class, and are then used to train the shallow random forest classifier. Once trained, this classifier can then be used to generate a segmentation (dense pixel classification, see **Fig. 1D**).

Two or more classes can be used to distinguish foreground objects from background pixels. **Fig. 2A & B** showcase examples of a single foreground and background classes. If desired, out of focus objects can even be discarded, for example by making such pixels part of the background class (**Fig. 2B**, arrowheads). For more complex segmentation tasks that need to discriminate various visible structures (*e*.*g*. nucleus vs. cytoplasm vs. background) or cell types (as in **Fig. 2C**), two or more foreground classes can be used (**Fig. 2D**).

**Fig. 2.**
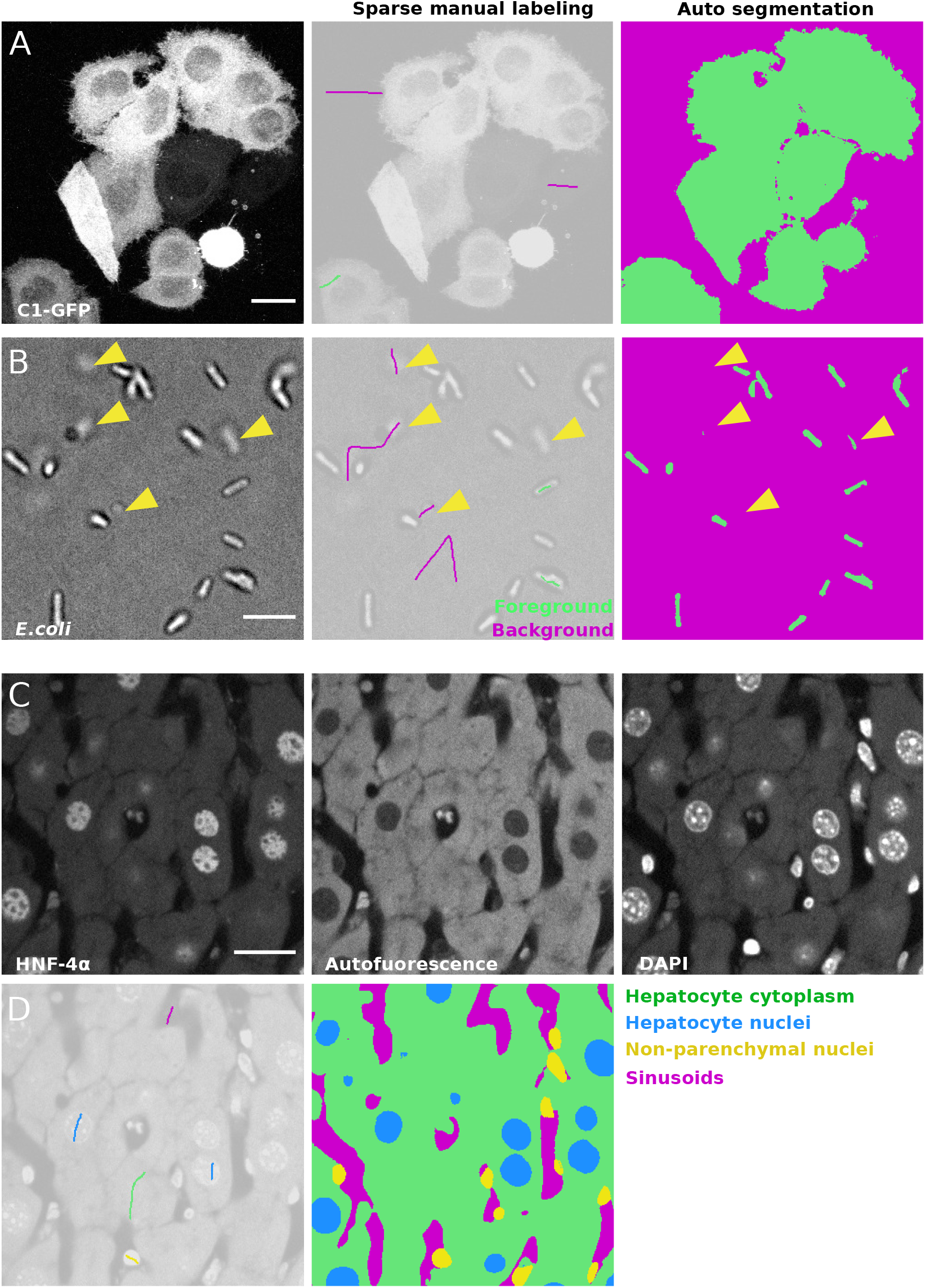
Semantic segmentation of microscopy images with Labkit’s pixel classification: **(A)** Maximum intensity projection of a confocal stack showing HeLa cells expressing C1-GFP (left), next to the sparse labeling (scribbles, center) and resulting cell segmentation (right). **(B)** Bright field microscopy image of *E*.*coli*, sparse labeling discriminating cells and background and the resulting segmentation. Arrowheads show that segmentation of out of focus objects can be reduced by including pixels of such objects in the background class. **(C)** Fixed mouse liver tissue section stained with immunofluorescence and imaged in multiple channels with a spinning disk confocal microscope, showing Hepatocyte nuclei stained with antibody against HNF-4*α* a transcription factor expressed in hepatocytes, hepatocyte cytoplasm (autofluorescence) and all nuclei stained with DAPI. **(D)** Labeling and resulting segmentation of the liver tissue section shown in A, segmenting Hepatocyte cytoplasm (green), Hepatocyte nuclei (blue), nuclei of non-parenchymal cells (yellow) and sinusoids (magenta). Scale bars 20 μm **(A), (C)** and 5 μm **(B)**

As opposed to deep learning algorithms, random forests are typically trained in a matter of seconds. Drawing scribbles and computing the segmentation can therefore conveniently be iterated due to the efficient parallelization we have implemented, leading to live segmentation. Live results are computed and displayed only on the currently visualized image slice in BigDataViewer to increase the interactivity. Hence, the effect of additional scribbles (sparse labels) is instantly visible and users can stop once the automated output of the pixel classifier reaches a similar quality to that of a fully manual pixel annotation. This iterative workflow makes working with Labkit very efficient, even when truly large image data are being processed. BigDataViewer’s bookmarking feature can additionally be used to quickly jump between previously defined image regions, thereby allowing validating the quality of the pixel classifier on multiple areas. Since we use ImgLib’s caching infrastructure, all image blocks that have once been computed are kept in memory and switching between bookmarks or browsing between parts of a huge volume is fast and visually pleasing. Once sufficiently trained, the classifier can be saved for later use in interactive Labkit sessions or in Fiji/ImageJ macros. The entire dataset can be directly segmented and the results saved to the disk. Recently, sparse labeling combined with random forest pixel classification in Labkit was used to segment mice epidermal cells (32), as well as mRNA foci in neurons (33).

Once the image is fully segmented, the generated segmentation masks can be transferred to label layers and the drawing tools can now be used to curate them. The goal of curation is to resolve the remaining errors made by the trained pixel classifier, such as drawing missing parts, filling holes, erasing mislabeling and deleting spurious blobs (**Fig. 3**). Label curation is performed until the curated segmentation is deemed satisfactory for downstream processing or analysis. Labkit can also be used to curate segmentation results obtained by other methods that are not available within Labkit, including deep learning based methods (34).

**Fig. 3.**
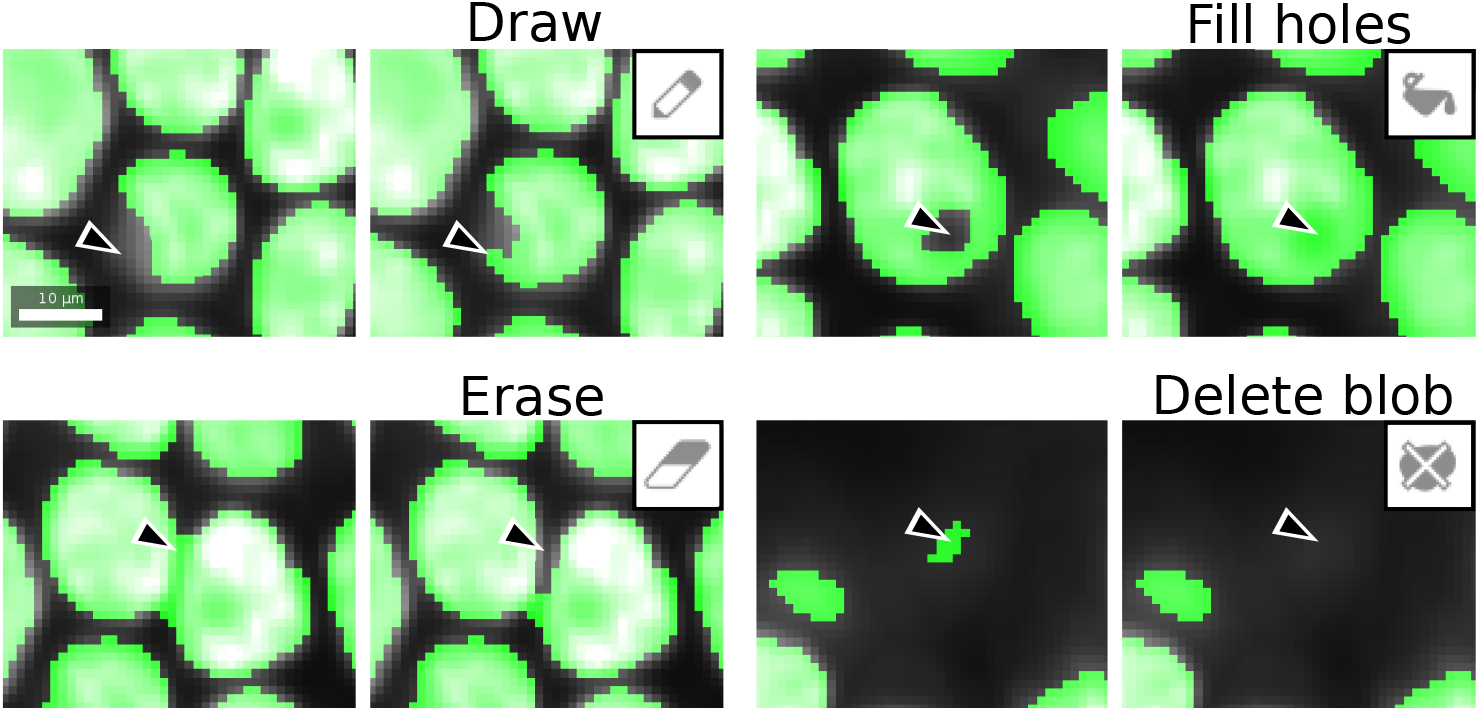
Labkit labeling tools used for curation: labels generated by manual dense labeling or automatic segmentation can be efficiently curated with drawing, filling, erasing, or deletion of entire objects. Scale bar 10 μm.

Automated segmentation with Labkit and the possibility to quickly curate any automated segmentation result make Labkit a powerful tool that can considerably shorten the time required to generate ground-truth data for training deep learning approaches. For example, we compared automatic and manual segmentation with Labkit on a rather small subset of images (N=26, see one example in **Fig. 4A**) made publicly available by the 218 Data Science Bowl (35). We segmented all images within 5 minutes by iterative scribbling and automated segmentation (see **Fig. 4B**). While many images consisted of homogeneous nuclei and led to high quality results, images with heterogeneous nuclei resulted in segmentation errors (see arrows in **Fig. 4B**). Such errors include spurious instances that do not correlate with any object in the original image, instances that correspond to the fusion of multiple instances, instances with holes, or even instances that split in two. Such errors are obviously undesirable and negatively impact the overall average precision score (AP = 0.72, see Methods for the metrics definition). As described above, all such segmentation errors can easily be corrected within Labkit, either by adding sparse labels corresponding to typical areas with errors, done during the iterative process, or when they persist by manually curating the residual errors in the final automated results (**Fig. 4C**). Curating all 26 images took an additional 10 minutes and raised the corresponding average precision to 0.76, a score very close to the inter-observer distance (AP = 0.78), as shown in **Fig. 4C & D**. In contrast, manually segmenting all images required more than an hour (**Fig. 4D**), which is four times longer than scribble-based pixel classification with Labkit, followed by full curation of the results to obtain images of comparable quality.

**Fig. 4.**
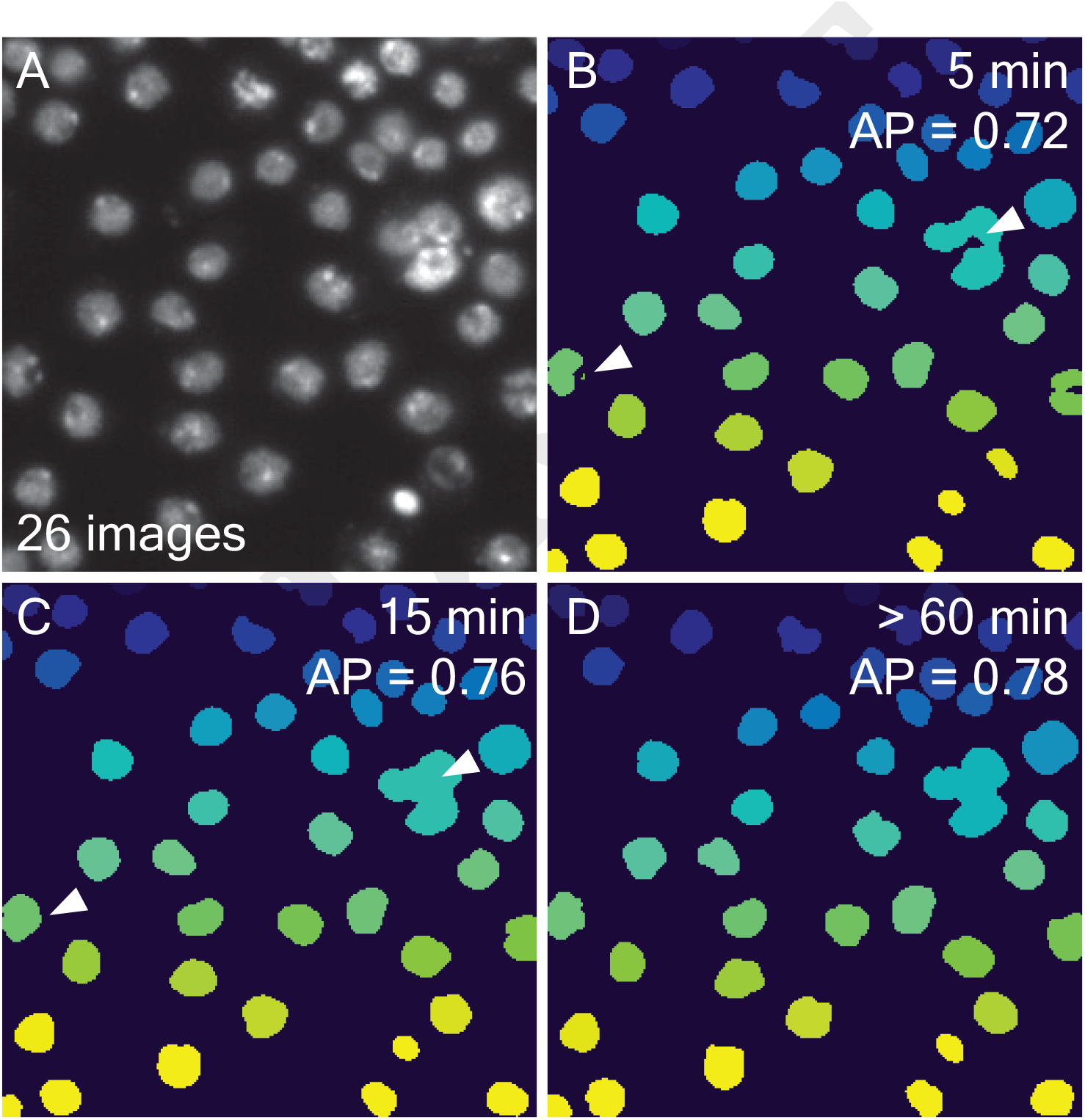
Comparing automatic and manual ground-truth generation with Labkit: **(A)** Fluorescence image of nuclei (out of 26 images) extracted from the 2018 Data Science Bowl (35). **(B)** Results from Labkit automated segmentation of **(A)** after extracting connected components and giving each instance a unique pixel value. The arrows point to various segmentation errors. On the top right corner, the total time necessary to obtain the corresponding segmentation of all 26 images (including labeling) is indicated. Below the timing is the average precision (see Methods) as compared to a dense manual labeling performed by another observer. **(C)** Curation of **(B)** with same post-processing. The arrows point to the corrected errors mentioned in **(B)**. The timing information includes **(B). (C)** Dense manual labeling of **(A)** and the same post-processing as in **(B)**. No scale bar was available for the images.

Hence, whenever Labkit automated segmentation is by itself not sufficient, manually curating the results yields ground-truth data that can be used to train a deep learning method, leading to higher segmentation quality with less labeling effort.

### Software and workflow integration

Labkit’s automatic segmentation is not limited to the dataset it was trained on. Because the trained classifier can be saved for later use, it can be applied to new images. While ensuring reproducibility of the results, it also helps maintaining consistency in the image segmentation. Manually loading both images and trained classifier in Labkit for multiple sets of images is a repetitive task ill-suited for automated workflow. Therefore, to simplify the integration into existing workflows in Fiji, Labkit can be easily called from the ImageJ macro language. For instance, a simple macro script can open multiple datasets and segment each of them using a trained classifier.

Image segmentation can be further accelerated by running the process on GPUs thanks to CLIJ (27). Once CLIJ properly set up, GPU acceleration is available for Labkit in both graphical interface and macro commands. GPU processing is particularly beneficial in the case of large images, for which it allows shortening the lengthy segmentation tasks. Performing GPU-accelerated segmentation in Labkit is a matter of activating a checkbox, and does not present additional complexity to users.

Some images, however, are far too large to be processed on a consumer machine in a reasonable amount of time, if they can be stored at all on such a computer. For such data, modern workflows resort to the use of HPC clusters, which are purposely built for high computing performances with large available memory. Labkit offers a command line tool (36) allowing advanced users to segment images on HPC clusters. The capability of extending Labkit and re-using its components is illustrated by integration with the commercial Imaris software (Oxford Instruments, UK) via the recently released ImgLib2-Imaris compatibility bridge. In this context, Labkit operates directly on datasets that are transparently shared (without duplication) between Imaris and ImgLib2 (25). These datasets can be arbitrarily large, as both Imaris and ImgLib2 implement sophisticated caching schemes. In the same fashion, output segmentation masks are transparently shared with the running Imaris application, making additional file import/export steps unnecessary. Importantly, this functionality can also be triggered and controlled directly from Imaris to integrate it into streamlined object segmentation workflows.

### Performance of Labkit

In order to process large images on consumer computers, software packages must be able to load the data in memory, process it and save the results, all within the constraints of the machine. In Labkit, this is achieved by reading only the portions of the image that are displayed to the user, thanks to the use of the HDF5 format (37) and the BigDataViewer (26). The image is further processed in chunks using a new ImgLib2 (25) implementation of the Trainable Weka Segmentation algorithm. As a result, Labkit is capable of processing arbitrarily large images and is compatible with GPU acceleration and distributed computation on HPC clusters.

To illustrate this, we segmented a 13.4 gigapixel image (482×935×495×60 pixels, 25 GB) on a single laptop computer, with and without GPU, and with different nodes of an HPC cluster (see **Table 1**). The image was extracted and 2x down-sampled from the *Fluo-N3DL-TRIF* dataset made available for the Cell Tracking Challenge (34, 38, 39) benchmark competition. Running the segmentation on the laptop using GPU acceleration sped up the computation by 7.5 fold, illustrating the benefit of harnessing GPU power for processing large images. While running computation on an HPC cluster comes with overhead, increasing the number of CPU nodes shortens the computation dramatically, reaching a 40fold improvement from 1 CPU node to 50. Finally, GPU nodes on an HPC allow for more parallelization of the computation and therefore even higher computational speed-up on the segmentation task, with 10 GPU nodes processing the data in slightly over a minute.

**Table 1.**
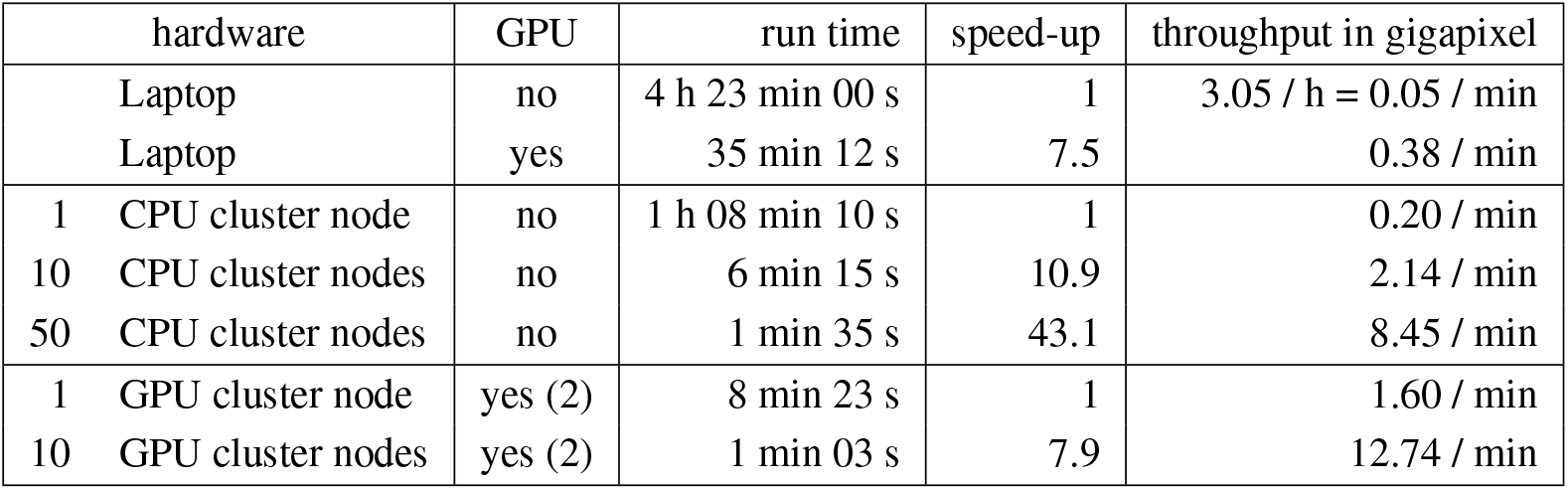
Benchmarking computation speed while segmenting a large biological image on various hardware: the experiment was performed on a laptop without and with GPU acceleration, and on different numbers of CPU and GPU cluster nodes. In each category, the speed-up is calculated in comparison to the slower entry. Numbers in between parenthesis in the GPU column indicate the number of GPU per cluster node.

| hardware | GPU | run time | speed-up | throughput in gigapixel |
| --- | --- | --- | --- | --- |
| Laptop | no | 4 h 23 min 00 s | 1 | 3.05 / h = 0.05 / min |
| Laptop | yes | 35 min 12 s | 7.5 | 0.38 / min |
| 1 CPU cluster node | no | 1 h 08 min 10 s | 1 | 0.20 / min |
| 10 CPU cluster nodes | no | 6 min 15 s | 10.9 | 2.14 / min |
| 50 CPU cluster nodes | no | 1 min 35 s | 43.1 | 8.45 / min |
| 1 GPU cluster node | yes (2) | 8 min 23 s | 1 | 1.60 / min |
| 10 GPU cluster nodes | yes (2) | 1 min 03 s | 7.9 | 12.74 / min |

Furthermore, we trained and optimized a classifier on the *Fluo-N3DL-TRIF* dataset (original sampling), the largest dataset of the Cell Tracking Challenge (training dataset of size 320 GB, evaluation dataset of size 467 GB), and submitted it for evaluation against undisclosed ground-truth. The segmentation of both training and evaluation datasets was performed on an HPC cluster. Labkit pixel classification ranked as the highest performing segmentation method on this dataset for all three evaluation metrics (*OP*_*CSB*_, *SEG* and *DET*) (40). More specifically, Labkit segmentation obtained the following scores: *OP*_*CSB*_ = 0.895 (0.886 for the second highest scoring entry), *SEG* = 0.793 (0.776) and *DET* = 0.997 (0.997), performing better than the other entries, including classical (bandpass segmentation) or deep learning (convolution neural network) algorithms. As opposed to the deep learning algorithms to which it was compared, Labkit only made use of a few hundred pixels in total, distributed throughout a small fraction of the training dataset (7 frames). Finally, Labkit’s classifier was simply trained through the Labkit graphical interface, illustrating its ease of use.

## Discussion and conclusion

Labkit is a labeling tool designed to be intuitive and simple to use. It features a robust pixel classification algorithm aimed at segmenting images between multiple classes with very little annotation required. Similar to other tools of the BigDataViewer family (26, 41–43), it integrates seamlessly into the SciJava and Fiji ecosystem. It can be easily installed through the Fiji updater and incorporated into established workflows using ImageJ’s macro language.The results of Labkit’s segmentation can be further analysed in Fiji or exported to other software platforms, such as CellProfiler (18), QuPath (20) or Ilastik (19).

Manual labeling, in both 2D and 3D, is also made easy by Labkit. Other alternatives exist, among which QuPath (20) (2D), Ilastik (19), napari (44) or Paintera (45). In particular, Paintera is specifically tailored to 3D labeling of crowded environment, but at the cost of a steeper learning curve.

Labkit is compatible with a wide range of image formats since image data can be loaded directly from Fiji using BioFormats (46). Nonetheless, in order to fully benefit from Labkit optimizations for large images, users must first convert their terabyte-sized images to a file format allowing highspeed access to arbitrary located sub-regions of the image. This strategy is also employed by other software, with the example of Ilastik (19). One such format is HDF5 (37), and Labkit uses in particular the BigDataViewer HDF5+XML version. In Fiji, images can easily be saved in this format using BigStitcher (42) or Multiview-Reconstruction (47, 48). In the Cell Tracking Challenge (39, 40), Labkit segmentation outperformed other entries on a particular dataset, among which two deep learning approaches. These methods were designed as part of a cell segmentation and tracking pipeline on various images, and it is likely that recent and more specialized segmentation algorithms, such as StarDist (11) or CellPose (15), would perform overall better. Yet, the full potential of deep learning algorithms is only reached when a sufficient amount of ground-truth data is available, which is too frequently the limiting factor. Generating ground-truth data for a deep learning method is a tedious endeavour without the insurance of a perfect segmentation result. A safer strategy is therefore to first try shallow learning for segmentation tasks, before even thinking of moving to deep learning algorithms. In cases where higher segmentation quality is truly necessary, curated results from shallow learning can be used to generate the massive amount of ground-truth required to train a deep learning algorithm. As seen previously, Labkit is useful in all these scenarios since it can be used to manually generate ground-truth annotations or to segment the images with shallow learning before curating the results in order to use them as ground-truth for other learning-based algorithms (see **Fig. 5**).

**Fig. 5.**
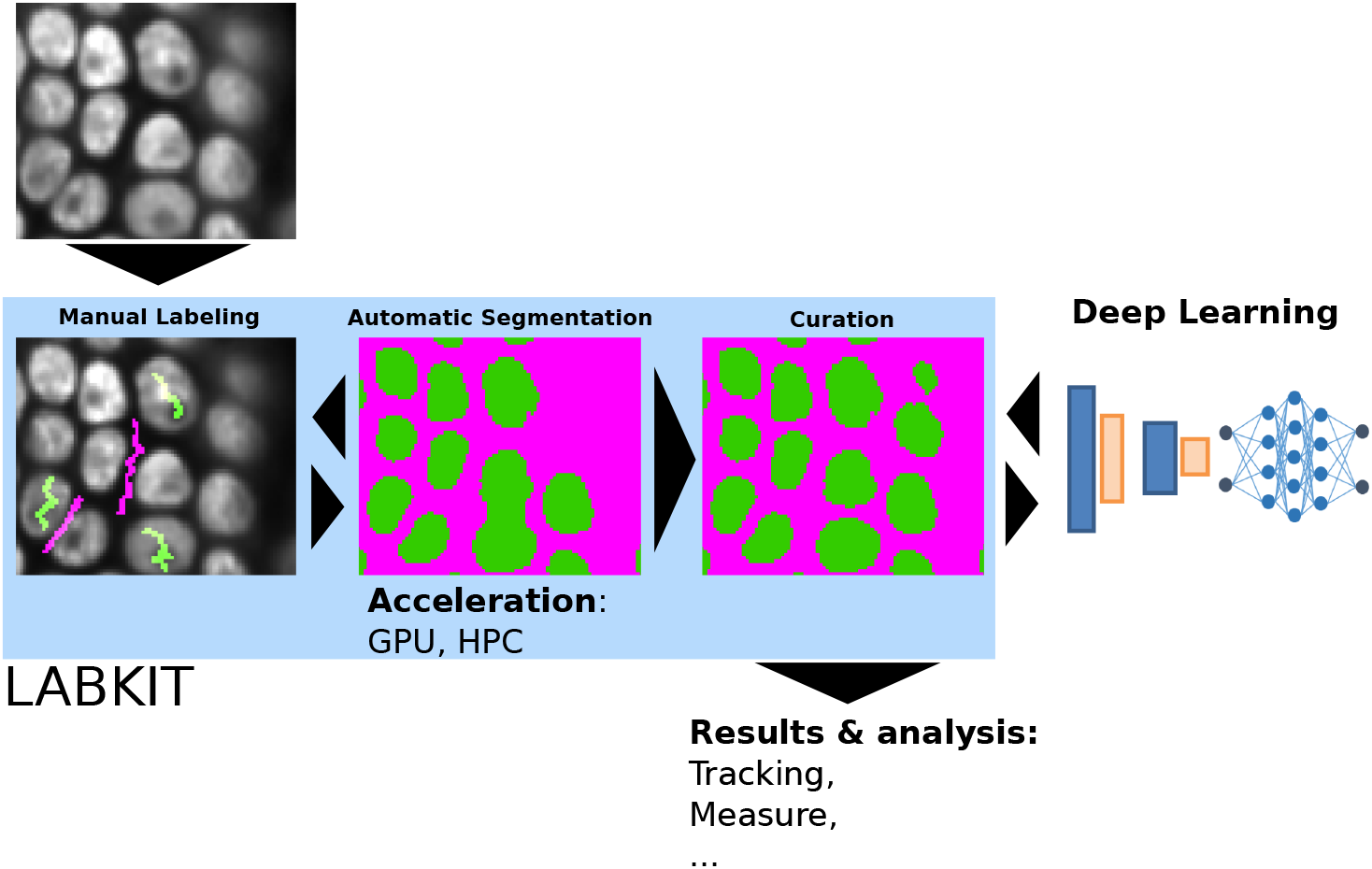
Labkit’s iterative and interactive segmentation used for ground-truth generation: manual labeling, automatic segmentation and curation in Labkit enable easy and rapid image segmentation, whose results can be further processed or used as ground-truth for deep-learning classifiers

In the future, we intend to extend Labkit’s functionalities to improve manual and automated segmentation. For instance, we will add a magic wand tool to select, fill, fuse or delete labels based on the pixel classification. Furthermore, we aim to add new pixel classifiers,such as the deep learning algorithm DenoiSeg (12) already available in Fiji. Labkit source code is open source and can be found online (49), together with its command-line interface (36) and tutorials (50).

## Methods

### A. Timing instance segmentation generation

The dataset consisted of all 256×256 images (N=26) in the test sample of StarDist (11), originally published as part of the 2018 Data Science Bowl (35) (subset of *stage1_train*, accession number BBBC038, Broad Bioimage Benchmark Collection). The images were loaded in Labkit as a stack and sparsely labeled (scribbles). A classifier was then trained with the default filter settings: “original image”, “Gaussian blur”, “difference of Gaussians”, “Gaussian gradient magnitude”, “Laplacian of Gaussian” and “Hessian eigenvalues”, with sigmas: 1, 2, 4 and 8. The results were saved and then manually curated using the brush and eraser tools. Finally, the same original image stack was densely manually labeled afresh. The total time required to process all images was measured using a chronometer for i) Labkit automated segmentation, including the sparse manual labeling, ii) the previous step followed by a curation step and iii) dense manual labeling. In order to evaluate the segmented images, connected components were computed (4-connectivity) and given unique pixel values (instance segmentation). Quality metrics scores were calculated as the average precision with threshold 0.5 as defined in StarDist (11). We used dense manual labeling performed by another observer as reference images, and computed the metrics score for the results obtained in i), ii) and iii). The average metrics over the images were calculated as a weighted average of each individual image, where the weights were the number of instances in the reference image.

### B. Speed benchmark

The dataset was downloaded from the Cell Tracking Challenge (39) website, and consisted of the first training dataset of the *Fluo-N3DL-TRIF* example. The dataset was down-sampled by a factor 2 in order to reduce its size and simplify the benchmarking. The dataset was then saved in the BigDataViewer XML+HDF5 format using BigStitcher (42). Labkit was used to draw a few scribbles on both background and nuclei areas, and to train a random forest classifier using the default settings. The trained model was then saved. The Labkit command line tool was used to run the benchmark experiment on a Dell XPS 15 laptop (32 MB RAM, Intel Core i7-6700HQ CPU with 8 cores, GeForce GTX 960M GPU) and on an HPC cluster, with both CPU (256 GB RAM, Intel Xeon CPU E5-2680 v3 with 2.5 GHz and 24 cores) and GPU (512 GB RAM, Intel Xeon CPU E5-2698 v4 with 2.2 GHz and 40 cores, with two GeForce GTX 1080 GPUs) nodes. The segmentation results on the HPC were saved in the N5 format to maximize writing speed. Benchmarking included read/write of image data form disc, optional data transfer to the GPU, computation of feature images and classification all together.

### C. Cell tracking challenge

As in the speed benchmark sample, all *Fluo-N3DL-TRIF* datasets (training and evaluation) were converted to BigDataViewer XML+HDF5 format using the BigStitcher Fiji plugin. This time, however, no down-sampling was applied to the images. For training, only frames 0, 1, 10, 20, 40, 50 and 59 from sequence “01” of the training dataset were used. A few hundred pixels were annotated as foreground and background. Only nuclei’s central pixels were labeled as foreground in order to force the classification algorithm to return segments of smaller size than the actual nuclei. Thus, segmented nuclei are unlikely to touch and segmentation errors are minimized. We used the following filters to train the random forest classifier: “original image”, “Gaussian blur”, “Laplacian of Gaussian”, and “Hessian eigenvalues”, with sigma values 1, 2, 4, 8 and 16. The filters can be set in Labkit’s interface through the parameters menu of the classifier. The trained classifier was saved and the evaluation dataset was segmented using the Labkit command line tool on an HPC. Since the output of the pixel classification is a binary mask, we performed a connected component analysis to assign unique pixel values to the individual segments. Finally, we dilated the segments to match the size of the nuclei. The dilation was done in three steps: the first two steps with a three-dimensional 6-neighborhood dilation kernel, then with a 3×3×3 pixel cube kernel. The combination of dilation kernels was chosen as to optimize the SEG score on the training dataset. All metrics scores were computed by the Cell Tracking Challenge platform.

## Author Contributions

F.J., M.A., T.P. and D.S. designed the project; M.A. implemented the software with help from T.P., D.S. and R.H.; M.A. and J.D. performed experiments; M.A., J.D., C.S. and F.J. wrote the manuscript with inputs from all authors.

## Data Availability Statement

Publicly available datasets were analyzed in this study. This data can be found here: https://bbbc.broadinstitute.org/BBBC038, accession number BBBC038, Broad Bioimage Benchmark Collection and http://celltrackingchallenge.net/3d-datasets/, Fluo-N3DL-TRIF dataset, Cell Tracking Challenge.

## ACKNOWLEDGEMENTS

We thank Anne Wuttke (Zerial lab, MPI-CBG), Sascha Kuhn (Nadler lab, MPI-CBG), Maria Luisa Romero Romero (Toth-Petroczy lab, MPI-CBG), Akanksha Jain & Anastasios (Tassos) Pavlopoulos (Tomancak, MPI-CBG) for sharing the experimental data. We also want to thank the Scientific Computing Facility at MPI-CBG for giving us access to HPC infrastructure.

